# Folate receptor overexpression shortens *C. elegans* lifespan by impeding adaptation to microbial metabolism

**DOI:** 10.1101/2023.06.26.546484

**Authors:** Bideep Shrestha, Milla Tallila, Olli Matilainen

## Abstract

Folate receptor (FR) alpha and beta (FRα and FRβ) are membrane-anchored transporters that mediate folate uptake through endocytosis. Unlike other folate transporters, FRs are expressed at low levels in normal tissues, while their expression is strongly increased in several cancers. In addition to its canonical role in folate transport, FRα, the most studied FR, also regulates signaling pathways unrelated to folate- or one-carbon metabolism. Nevertheless, it is not known how loss of folate receptors, or their overexpression, affects health- and lifespan. To elucidate this, we utilized *Caenorhabditis elegans*, in which FOLR-1 is the sole homolog of folate receptors. Interestingly, loss of FOLR-1 does not affect reproduction, fitness, proteostasis or lifespan, indicating that it is not required for folate transport to maintain health. Strikingly, in contrast to *folr-1* depletion, *folr-1* overexpression shortens lifespan in a cocultured *E. coli* strain-dependent manner. Furthermore, we found that *folr-1* overexpression blunts the lifespan extension upon treatment with sulfamethoxazole, a sulfonamide that promotes longevity by limiting folate in *E. coli*. These data suggest that animals overexpressing *folr-1* are deficient in their ability to adapt to changes in microbial metabolism, thus revealing an intriguing and non-canonical role of FR in lifespan regulation. Therefore, this work could serve as a basis for further studies to elucidate the organismal effects of abnormal FR expression in diseases such as cancer.

**Author Summary:** Folate receptors (FRs) differ from other folate transporters based on, for example, these two things: their expression is strongly restricted, and they can regulate intracellular cellular processes independently of folate- and one-carbon metabolism (OCM). Interestingly, FRs have an essential role in embryonic development, but on the other hand, their overexpression in cancer have been shown to reduce patient survival. Apart from development and disease, the role of FRs in aging remains unknown. By utilizing *C. elegans*, we show that FOLR-1 (FR homolog in *C. elegans*) is not required to maintain normal physiology, whereas its overexpression shortens lifespan through a mechanism dependent on cocultured bacteria. Since cocultured bacteria constitute the *C. elegans* gut microbiota, this study places elevated FR expression as a link between gut commensal bacteria and organism’s lifespan, raising the intriguing question of whether the same mechanism applies in cancer.

## Introduction

Folate, also known as vitamin B_9_, is a cofactor in one-carbon metabolism (OCM), which supports processes such as purine and thymidine biosynthesis, homeostasis of glycine, serine, and methionine, epigenetic maintenance, and redox defense [1]. Eukaryotic organisms such as nematodes, flies and mammals cannot synthesize folates, and therefore, they must obtain it from the diet or through biosynthesis within the gut microbiota. Due to its central role in OCM, folate deficiency contributes to multiple pathologies including cancer, cardiovascular disease, and developmental anomalies such as neural tube defects [2]. In addition to their role in preventing diseases and promoting development, folate and the associated OCM regulate aging in multiple organisms [3]. As an examples from *C. elegans*, it has been shown that altered function of OCM is a common signature of long-lived *C. elegans* strains, and that OCM downregulation triggers lifespan-extending methionine restriction [4]. Moreover, metformin, a drug to treat type-2-diabetes, inhibits microbial folate and methionine metabolism, which leads to extended lifespan through methionine restriction [5]. Moreover, it has been shown that inhibition of microbial folate synthesis leads to extended longevity [6, 7], which further strengthens the connection between folate, microbiome, and aging. Hence, based on extensive research on folate and its metabolism, it can be said that this vitamin plays a key role in every stage of the organism’s life.

In humans, majority of folate is transported to cells by three transporters: proton-coupled folate transporter (PCFT), reduced folate carrier (RFC) and folate receptors (transport folates into cells via an endocytic mechanism). PCFT functions mainly in the upper gastrointestinal tract [8], whereas RFC is ubiquitously expressed [9]. Although RFC is the major factor transporting folate to tissues, it binds folates with relatively low affinity (K_m_ = 1–10 μM) [10]. In contrast to RFC, folate receptors (FRs, include three folate-binding isoforms: alpha (FRα), beta (FRβ) and gamma (FRγ)) bind folates with high affinity [11]. For example, FRα binds synthetic folic acid (Kd: <1 nM) and 5-methyltetrahydrofolate (5-MTHF) (Kd: 1–10 nM) efficiently at low physiologic concentrations [12]. Unlike RFC, folate receptors show restricted expression profile. The expression of constitutively secreted FRγ can be detected in normal and leukemic hematopoietic tissues, whereas the membrane-bound FRβ can be found in placenta and hematopoietic cells [13]. The membrane-bound FRα, the most studied and widely expressed FR, is expressed in choroid plexus, lung, thyroid, and kidney [13–16]. Notably, although FRs show low expression in normal cells, their expression is strongly increased in multiple cancers [14, 15, 17–21]. Even though the expression profiles of FRs’ have been well documented, the factors regulating their expression are still unknown.

*C. elegans* has two identified folate transporters, FOLT-1 (homolog of RFC) [22, 23] and FOLR-1 (homolog of FR) [24]. *folt-1* is ubiquitously expressed with pharynx and intestine displaying the strongest expression [22]. Loss of FOLT-1 causes germline and somatic defects, as well as shortened lifespan [23], thus demonstrating the importance of this folate transporter for *C. elegans* development and normal aging. Regarding FOLR-1, it has been shown that bacterial folates stimulate germ cell proliferation through this receptor [24]. This is an interesting observation because folates, which have a role as vitamins, do not stimulate germ cell proliferation, thus demonstrating that FOLR-1 is able to regulate germ cell number independently of OCM [24]. The OCM-independent role of FR is not limited to *C. elegans*, as multiple studies have shown that FRα is a regulator of JAK–STAT3 and ERK1/2 signaling pathways in mammalian systems [17]. Moreover, it has been shown that FRα is able to translocate into the nucleus and function as a transcription factor [25–27], providing further evidence for non-canonical functions of FRα.

As mentioned earlier, folate metabolism and OCM are important regulators of aging [4–7, 23]. Although FR can function both as a part of OCM and independently of it [17], it is not known whether FR modulates health and longevity. To address this, we created *C. elegans* lines lacking or overexpressing folate receptor *folr-1*. Interestingly, whereas loss of FOLR-1 does not affect health or longevity, we found that *folr-1* overexpression (OE) shortens lifespan in a *E. coli* strain-dependent manner. Furthermore, our data suggest that *folr-1* OE animals are unable to fully adapt to changes in microbial metabolism, which affects their lifespan. Together, these data shed light on the role of FOLR-1 in organismal physiology and reveal previously unknown linkage between this receptor and aging.

## Results

### FOLR-1 is expressed only in vulval cells, and the expression in other tissues is inhibited by histone-binding protein LIN-53

As the first step, we examined how *folr-1* is expressed in *C. elegans*. For this purpose, we used a strain in which endogenous FOLR-1 is tagged with fluorescent mNeonGreen-protein by CRISPR/Cas9-mediated genome editing. Interestingly, when imaging FOLR-1::mNeonGreen-strain at L4 larval stage, we found that FOLR-1 is expressed only in few vulval cells (Fig 1A). This vulval expression can also be seen in adult animals, but not in L3 stage (S1 Fig), demonstrating that FOLR-1 is expressed in a development-dependent manner. FOLR-1::mNeonGreen fusion protein forms circular shapes (Fig 1A and S1 Fig), suggesting that, like its mammalian homolog, it is located at the cell membrane. Due to its restricted expression pattern, it is possible that there are specific factors that inhibit FOLR-1 expression in other tissues. LIN-53, the homolog of mammalian histone-binding protein RbAp48, has been shown to repress the transcription of genes determining vulval cell fates [28]. Furthermore, it has been demonstrated that germ cells acquire the ability to be reprogrammed into distinct neuron types upon loss of *lin-53* [29]. Therefore, we asked whether LIN-53 regulates FOLR-1 expression. Strikingly, we found that *lin-53* RNAi induces ectopic FOLR-1 expression in intestine and gonad (Fig 1B), hence demonstrating that LIN-53, a factor that is required for muscle integrity and normal lifespan [30], controls the restricted expression pattern of FOLR-1.

**Fig 1.**
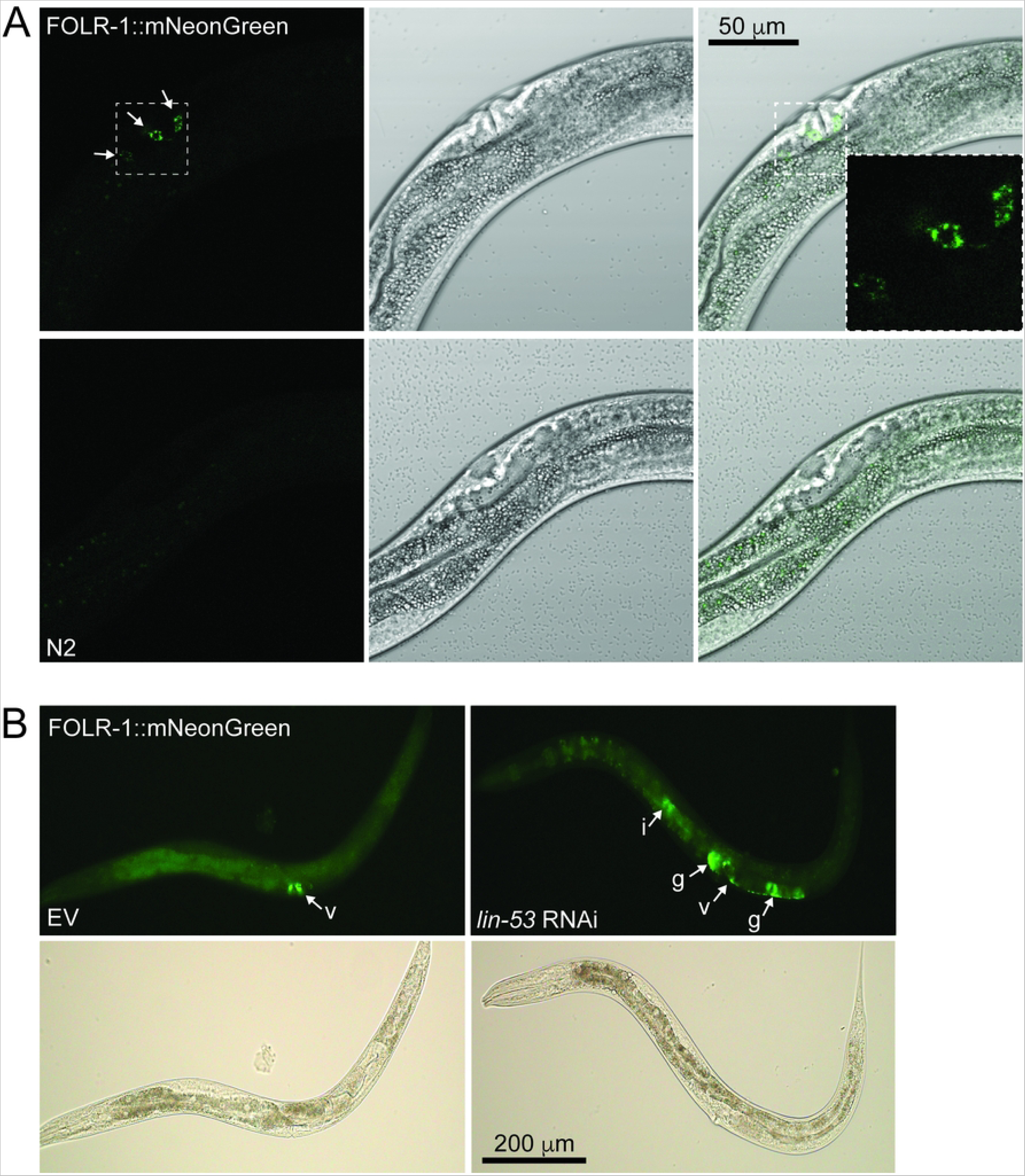
FOLR-1 expression profile. (A) Confocal images of L4 stage N2 (wild-type) and FOLR-1::mNeonGreen fusion protein-expressing animals. N2 is used as a control to recognize background signal. Arrows indicate FOLR-1::mNeonGreen localization. Dotted square marks the magnified area shown on the right. (B) Images of L4 stage FOLR-1::mNeonGreen animals treated with EV (empty vector) or *lin-53* RNAi. Arrows indicate FOLR-1::mNeonGreen expression (v, vulva; i, intestine; g, gonald).

### Loss of FOLR-1 does not affect lifespan, activity, progeny number, or proteotoxicity

Next, we asked whether loss of FOLR-1 influences animal physiology. For this purpose, we utilized two separate loss-of-function *folr-1* mutant strains generated by CRISPR/Cas9-mediated genome editing. The *folr-1(syb4116)* allele has a premature stop codon after 20 amino acids, whereas the *folr-1(syb5135)* allele is a complete deletion of *folr-1* gene (from start to stop codon) (Fig 2A). We found that both *folr-1* mutant alleles have normal lifespan (Fig 2B and Fig 2C). Additionally, loss of *folr-1* does not affect survival in higher temperature (S2A Fig). As a next step we investigated whether folate supplementation extends lifespan, and whether FOLR-1 is required for this process. Folic acid (FA) is a synthetic form of folate that is widely used as a food supplement. Interestingly, FRα has 14-fold higher affinity for FA compared to naturally occurring 5-methyltetrahydrofolate (5-MTHF) [31], and it has been published that 10 μM and 25 μM FA solutions spread on agar plates with *E. coli* OP50 as a food source extends *C. elegans* lifespan [32]. In our experiment we supplemented NGM agar with FA at final concentration of 10 μM and used *E. coli* HT115 as a food source. In these experiments, FA did not affect the lifespan of N2 or *folr-1(syb5135)* mutants (Fig 2D). Notably, in our experiments FA did not affect N2 lifespan even when used at final concentration of 250 μM (S2B Fig). Next, we examined the lifespan effects of 5-MTHF in N2 and *folr-1(syb5135)* animals. 5-MTHF is an intermediate in OCM, and like FA, it is also used as food supplement. Interestingly, since decreased OCM activity extends lifespan, 5-MTHF shortens the lifespan of OCM-deficient animals (and long-lived mutants) when used at low concentration (10 nM) [4]. Surprisingly, we found that 5-MTHF used at final concentration of 100 nM in NGM agar extends the lifespan of both N2 and *folr-1(syb5135)* animals (Fig 2E). These data demonstrate that 5-MTHF is a potent modulator of longevity, and that FOLR-1 is not required for the lifespan-extending effect of this folate species.

**Fig 2.**
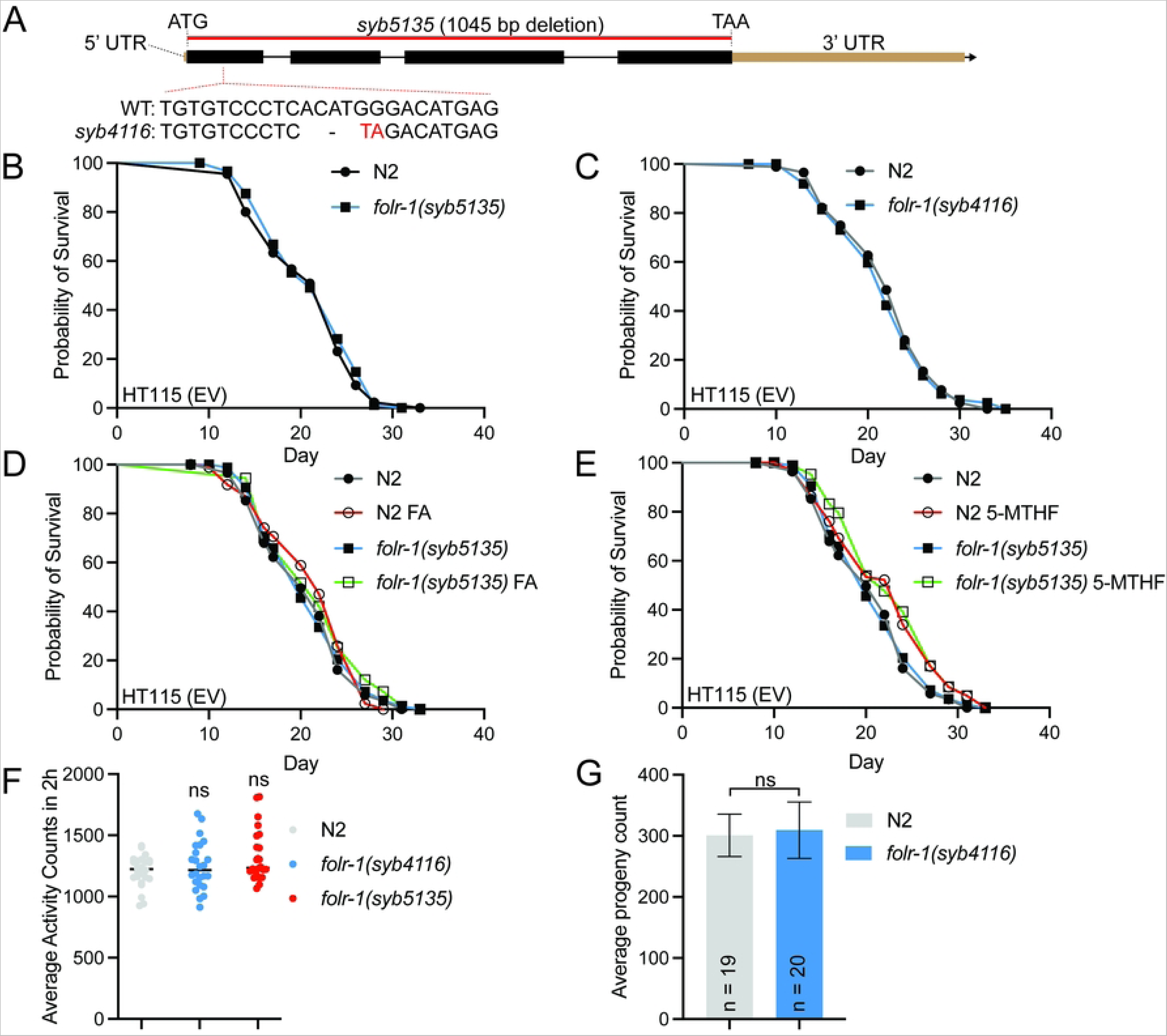
Loss of FOLR-1 does not affect lifespan, activity, or brood size. (A) Schematic presentation of *folr-1* gene with mutations generated for this study. (B) Lifespan of *folr-1(syb4116)* and (C) *folr-1(syb5135)* mutants on HT115 (carrying empty vector, EV) compared to N2. (D-E) Lifespan of *folr-1(syb5135)* mutant and N2 on plates supplemented with (D) 10 μM folic acid (FA) or (E) 100 nM 5-methyltetrahydrofolate (5-MTHF). Lifespan statistics are reported in S1 Table. (F) Activity of day 4 adult N2, *folr-1(syb4116)* and *folr-1(syb5135)* mutants measured with wMicroTracker. Each dot represents a group of 10 animals (n = 240 animals per condition). Data is combined from two independent experiments. Statistical significances were calculated with one-way ANOVA with Tukey’s test. (G) Brood size of N2 and *folr-1(syb4116)* mutants. Data is combined from two independent experiments. Statistical significance was calculated with unpaired Student’s *t*-test.

In addition to its role in longevity, we also investigated whether loss of FOLR-1 affects fitness. Firstly, we measured the activity of *folr-1* mutants at day 4 of adulthood. In line with the observations from lifespan assays (Fig 2B and Fig 2C), *folr-1* mutants do not show difference in activity compared to N2 animals (Fig 2F). Secondly, since loss of FOLT-1 causes sterility [23], we tested whether FOLR-1 affects reproductive fitness, and found that loss of this receptor does not affect the brood size (Fig 2G). These data support earlier study reporting that *folr-1* RNAi does not affect the number of eggs laid [24].

Folate deficiency increases the risk of Alzheimer’s disease (AD), whereas sufficient folate intake protects from this disorder [33]. Since the toxicity of amyloid beta (Aβ) oligomers are a central part of AD, we asked whether *folr-1* mutation modulates Aβ toxicity. For this purpose, we crossed *folr-1(syb4116)* mutant with GMC101, a strain expressing Aβ_1-42_ peptide in body-wall muscle cells [34]. Western blot experiment showed that *folr-1(syb4116)* mutation does not affect Aβ accumulation (Fig 3A). Furthermore, *folr-1* depletion does not affect the fitness of GMC101 animals (Fig 3B), demonstrating that FOLR-1 is not required to maintain proteostasis. Interestingly, in contrast to loss of FOLR-1, we found that *folt-1* RNAi increases the accumulation of Aβ oligomers (Fig 3C), which supports previous findings reporting that folate plays a protective role in AD [33].

**Fig 3.**
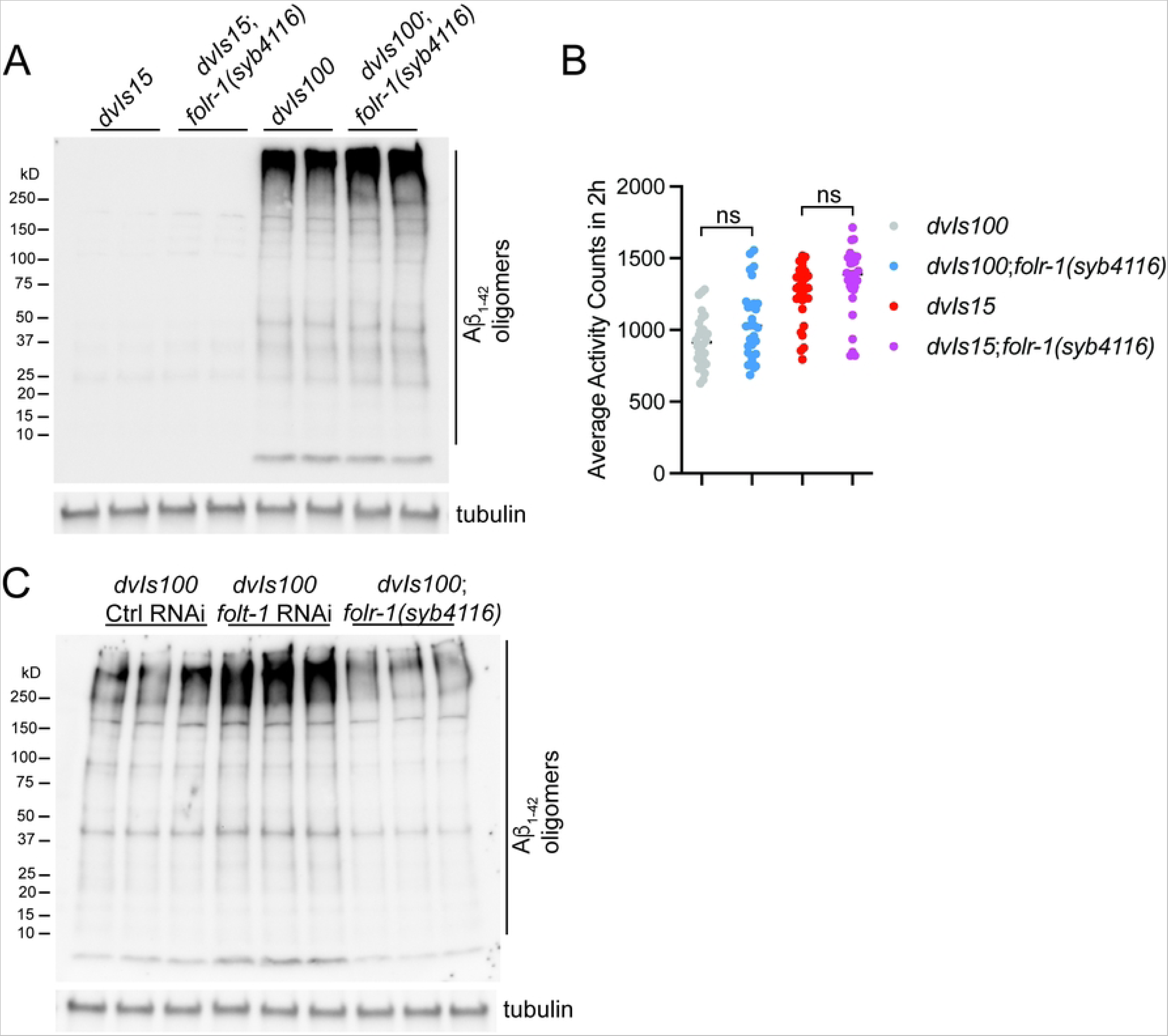
Loss of FOLR-1 does not aggravate proteotoxicity. (A) Amyloid beta (Aβ) Western blot and (B) activity of day 2 adult Aβ-expressing (GMC101, expresses transgene *dvIs100*)- and control strain (CL2122, expresses transgene *dvIs15*) in wild-type- and *folr-1(syb4116)* background measured with wMicroTracker. In (B), each dot represents a group of 10 animals (n = 300 animals per condition). Data is combined from three independent experiments (**p < 0.01, ****p < 0.0001, one-way ANOVA with Tukey’s test). (C) Aβ Western blot of Aβ-expressing strain upon control- and *folt-1* RNAi, and with *folr-1(syb4116)* background.

### Loss of FOLR-1 induces minor changes in ribosomal gene expression

Although loss of FOLR-1 does not induce any detectable phenotypes, we asked how it affects gene expression. For this purpose, we performed RNA-seq analysis to compare gene expression in L4 stage N2 and *folr-1(syb4116)* animals. Data analysis revealed that the expression of 451 and 447 genes are up- and downregulated, respectively, in *folr-1(syb4116)* animals (Fig 4A). When performing KEGG pathway [35] enrichment analysis for genes upregulated in *folr-1(syb4116)* mutants, we found that ribosome, the macromolecular machine mediating protein synthesis, shows the most significant enrichment (Fig 4B). On the other hand, downregulated genes in *folr-1(syb4116)* mutants did not show any significantly enriched KEGG pathway. To validate the RNA-seq data, we performed qRT-PCR analysis of selected ribosomal genes that were found to be significantly upregulated in *folr-1(syb4116)* mutants in RNA-seq. Interestingly, when comparing with the RNA-seq data, qRT-PCR did not show as clear upregulation of ribosomal genes, as the expression of many genes was found to be unchanged in *folr-1* mutants (S3A Fig and S3B Fig). For further validation, we used a strain in which endogenous RPS-6 is tagged with fluorescent wrmScarlet by CRISPR-Cas9-mediated genome editing. As expected, RPS-6 signal can be detected in every cell of L4 larvae, but we did not detect any differences between wild-type- and *folr-1(syb5135)* background (S3C Fig). Similarly to RPS-6::wrmScarlet imaging, RPS-6 Western blots from whole-animal extracts of day 2 adults did not reveal any differences between N2 and *folr-1(syb4116)* mutants (S3D Fig and S3E Fig).

**Fig 4.**
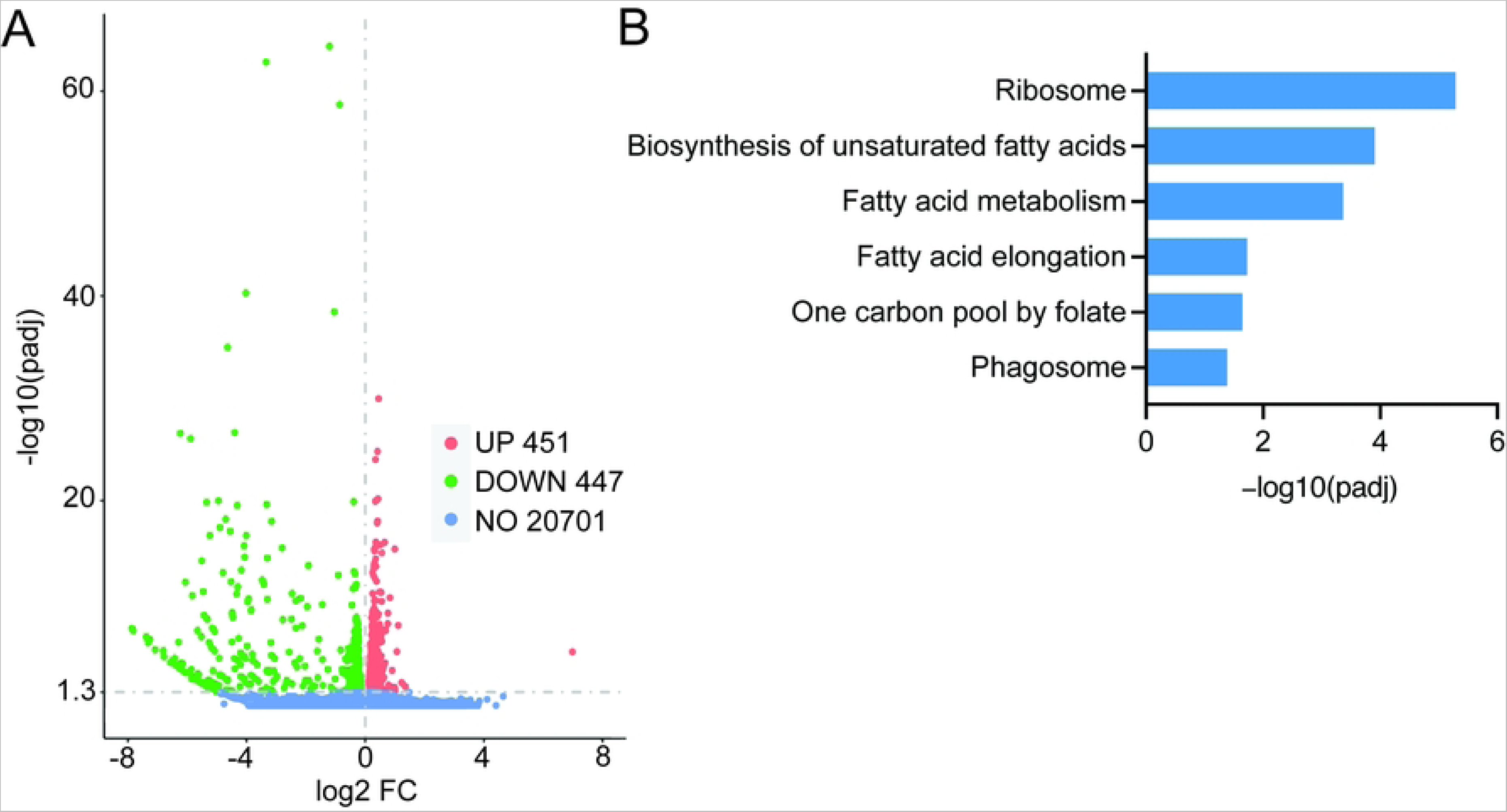
The effect of *folr-1* mutation on gene expression. (A) Volcano plot showing differentially expressed genes in L4 stage *folr-1(syb4116)* mutants compared to N2. (B) Enriched KEGG pathways among upregulated genes in *folr-1(syb4116)* mutants.

Together, the data presented above provide strong evidence that, unlike loss of folate transporter FOLT-1 [23], the depletion of *folr-1* has a minor effect on gene expression, and does not affect health- or lifespan in *C. elegans*.

### *folr-1* overexpression deteriorates fitness and shortens the lifespan of animals grown on *E coli* HT115

As mentioned earlier, FR is overexpressed in multiple cancers [14, 15, 17–21]. For example, the expression of FRα has been shown to be 100–300 times higher in breast, lung, kidney, and ovarian cancers when compared to healthy cells [18]. At protein level, the highest FRα expression has been detected in ovarian and brain carcinomas, in which FRα level can be up to 30 and 14 times higher compared to normal cells, respectively [14]. Since FRα overexpression in tumors has been associated with increased cancer progression and poor patient prognosis [17, 36], we asked how increased FR expression affects the physiology of *C. elegans*, in which all somatic cells in adult animal are post-mitotic. For this purpose, we used two independent *folr-1* overexpression (OE) strains (PHX4824 and PHX4825). These strains carry extra copies of the *folr-1* gene under its own promoter sequence. To ensure efficient expression of the transgene, *folr-1* 3’UTR was replaced with *unc-54* 3’UTR. Strikingly, we found that both *folr-1* OE strains have significantly shortened lifespan on *E. coli* HT115 (Fig 4A). To confirm that shortened lifespan is due to *folr-1* overexpression, we asked whether *folr-1* RNAi rescues the short lifespan of these strains. Before this experiment, we measured how efficiently *folr-1* RNAi downregulates *folr-1* mRNA level in *folr-1* OE strains. To ensure maximal knockdown efficiency, we placed L1 larvae of P0 generation on *folr-1* RNAi and extracted RNA from L4 larvae of the next generation (F1 generation), thereby exposing animals to *folr-1* RNAi for two generations. Interestingly, qRT-PCR analysis revealed that *folr-1* RNAi partially blunts *folr-1* overexpression in *folr-1* OE strain PHX4824, but not in *folr-1* OE strain PHX4825 (Fig 5B and Fig 5C). This may be because *folr-1* OE strain PHX4825 shows higher *folr-1* expression than PHX4824 strain when compared to N2 (Fig 5B and Fig 5C), and the high *folr-1* expression in PHX4825 strain may saturate the organismal RNAi capacity to downregulate this gene. For the lifespan analysis, we performed two independent lifespan experiments with *folr-1* RNAi. In first experiment, *folr-1* RNAi partially rescued the short lifespan of *folr-1* OE strain PHX4824 (Fig 5D), whereas in second experiment the lifespan of *folr-1* OE strain PHX4824 was fully rescued (see Table S1). However, according to the qRT-PCR data showing that *folr-1* RNAi does not reduce *folr-1* mRNA level in *folr-1* OE strain PHX4825 (Fig 5C), *folr-1* RNAi does not rescue the lifespan of this strain (Fig 5D and Table S1). Together, these experiments strengthen the conclusion that increased *folr-1* expression shortens lifespan.

**Fig 5.**
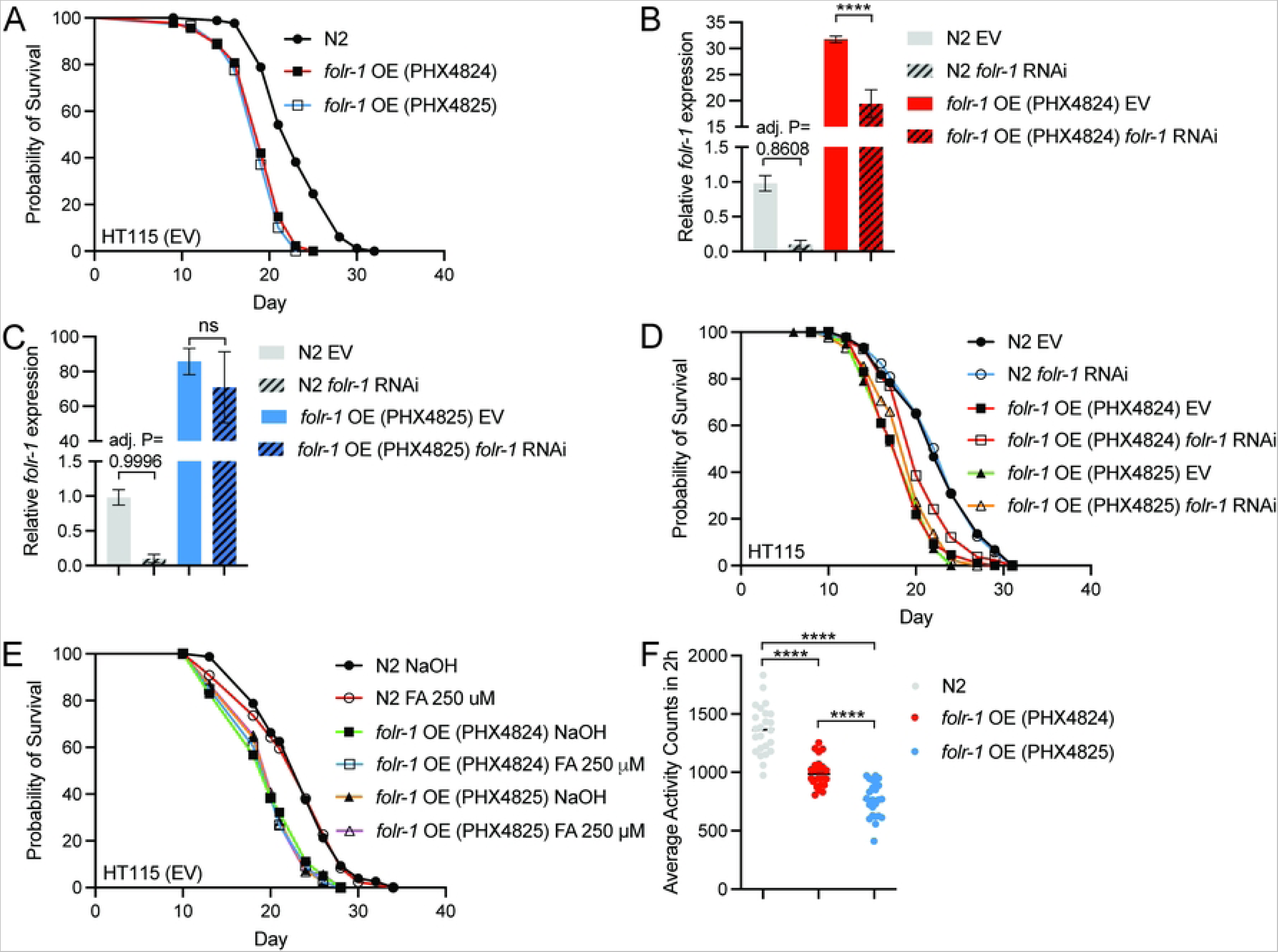
*folr-1* overexpression shortens lifespan and reduces activity on *E. coli* HT115. (A) Lifespan of *folr-1* OE strains (PHX4824 and PHX4825) on HT115 (EV) compared to N2. (B) *folr-1* qRT-PCR of *folr-1* overexpression (OE) strains PHX4824 and (C) PHX4825 upon control and *folr-1* RNAi. Data in (B) and (C) are from the same experiment divided into two graphs. Bars represent *folr-1* mRNA levels with error bars indicating mean ± SD of three biological replicates, each with three technical replicates (****p < 0.0001, one-way ANOVA with Tukey’s test). (D) Lifespan of N2 and *folr-1* OE strains on *folr-1* RNAi. (E) Lifespan of N2 and *folr-1* OE strains on plates supplemented with 250 μM folic acid (FA). Lifespan statistics are reported in S1 Table. (F) Activity of day 4 adult N2 and *folr-1* OE strains measured with wMicroTracker. Each dot represents a group of 10 animals (n = 240 animals per condition). Data is combined from two independent experiments (****p < 0.0001, one-way ANOVA with Tukey’s test).

Next, we asked whether *folr-1* OE strains have shortened lifespan due to excess absorption of folate. Therefore, we tested whether supplementing plates with folic acid leads to further decrease in their lifespan. However, folic acid supplementation does not affect the lifespan of *folr-1* OE strains (Fig 5E), indicating that their short lifespan is not due to possible folate toxicity. Finally, to test whether increased *folr-1* expression affects fitness, we measured the activity of *folr-1* OE strains at day 4 of adulthood. In contrast to *folr-1* mutants (Fig 2F), both strains show significant reduction in activity when compared to N2 (Fig 5F). These data are a striking demonstration of how FR overexpression can deteriorate animal’s health, thus providing possible explanation on why FR expression is tightly controlled in both *C. elegans* (Fig 1A, Fig 1B and S1 Fig) and humans [13–15].

### *folr-1* overexpression does not affect the lifespan of animals grown on *E. coli* OP50

*C. elegans* is maintained on a bacterial lawn, which also forms its gut microbiota. Importantly, it has been shown that changes in microbial metabolism have a significant impact on longevity [5–7, 37]. In this context, *E. coli* HT115 and OP50 are the two most commonly used bacterial strains in *C. elegans* experiments, and even these bacteria, which differ in their metabolism and transcriptome, have different effects on lifespan [38]. Since *C. elegans* has shortened lifespan when grown on OP50 compared to HT115 (on which the above-described lifespan experiments were performed) [5, 38], we asked how OP50 affects the longevity of *folr-1* mutants and OE strains. We found that N2, *folr-1(syb4116)* and *folr-1(syb5135)* mutants have similar lifespan on OP50, which is shorter than on HT115 (Fig 6A). Interestingly, although *folr-1* OE strains have shortened lifespan on HT115, their lifespan is similar to that of N2 and *folr-1* mutants on OP50 (Fig 6A), suggesting that *folr-1* OE strains are unable to adapt to healthier HT115 bacteria.

**Fig 6.**
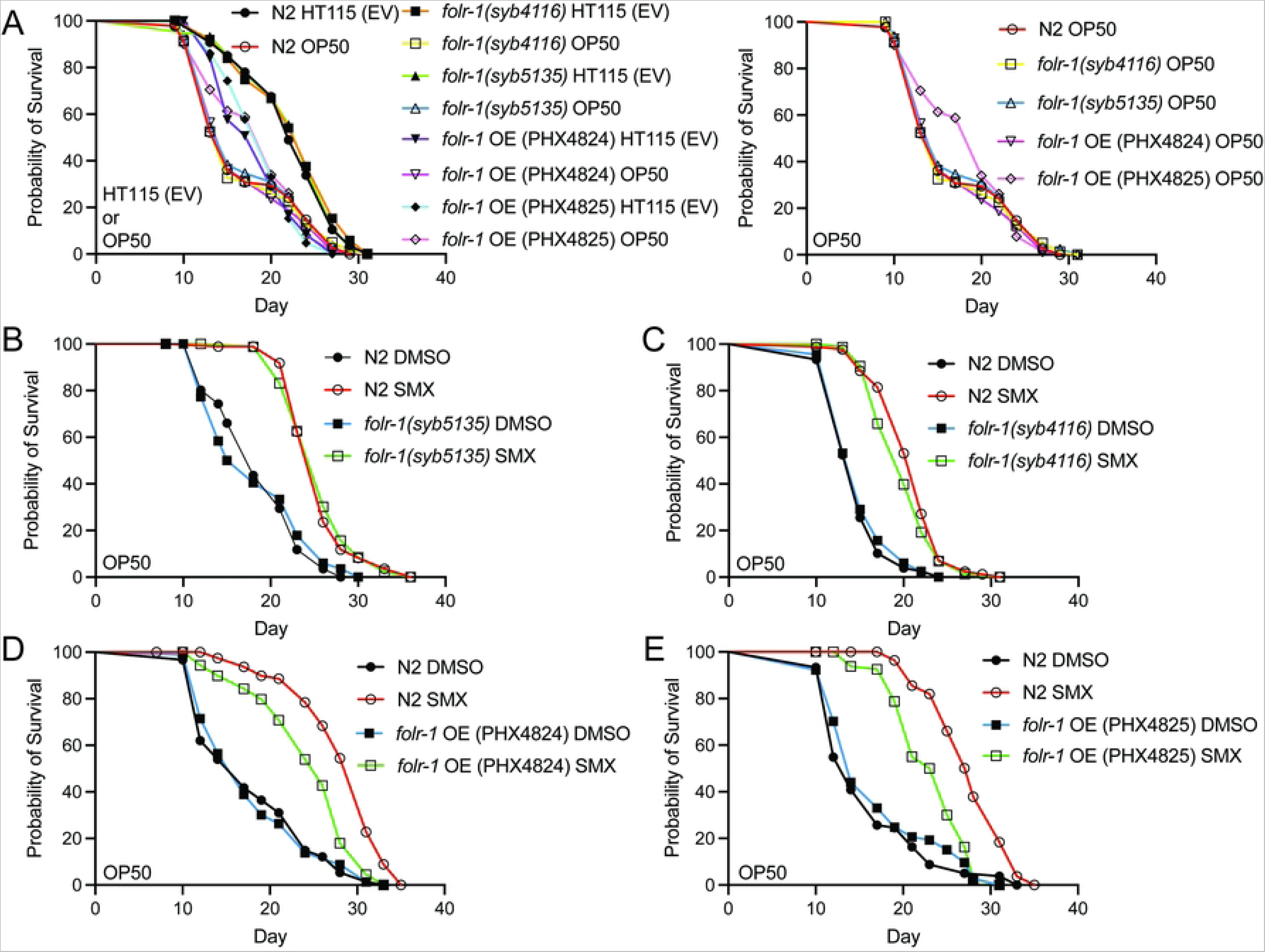
*folr-1* overexpression does not shorten the lifespan on *E. coli* OP50 but blunts SMX-mediated longevity. (A) Lifespan of N2, *folr-1* OE strains (PHX4824 and PHX4825) and *folr-1* mutants (*folr-1(syb4116)* and *folr-1(syb5135)*) on HT115 (EV) and OP50. The graph on the right shows the data for OP50-fed animals only. (B) Lifespan of N2, *folr-1(syb5135)* and (C) *folr-1(syb4116)* on SMX-supplemented OP50 plates. (D) Lifespan of N2, *folr-1* OE strain PHX4824 and (E) *folr-1* OE strain PHX4825 on SMX-supplemented OP50 plates. Lifespan statistics are reported in S1 Table.

Sulfamethoxazole (SMX) is a sulfonamide drug that extends *C. elegans* lifespan through inhibition of microbial folate synthesis [6, 7]. Importantly, the lifespan-extending effect of SMX is based on its ability to limit folate in *E. coli*, but not in *C. elegans* [7]. Therefore, we asked whether *folr-1* OE animals can adapt to SMX-induced changes in microbial metabolism. First, we found that loss of FOLR-1 does not affect the SMX-induced extension in lifespan (Fig 6B and Fig 6C). In contrast to *folr-1* mutants, we found that *folr-1* OE reduces the SMX-mediated longevity (Fig 6D and Fig 6E). Together with the finding that *folr-1* OE shortens lifespan on HT115 but not on OP50 (Fig 6A), these data demonstrate that animals overexpressing *folr-1* have a reduced ability to adapt to alterations in microbial metabolism, which then limits their lifespan in symbiosis with a more favorable microbiome.

## Discussion

Our work show that, unlike the depletion of RFC homolog FOLT-1 [23], loss of FOLR-1 does not affect health- or lifespan, thus supporting the statement by Chaudhari *et al*. that FOLR-1 is not essential for the uptake of folates to function as vitamins [24]. In contrast to *C. elegans*, FR ablation leads to embryonic lethality in mice [39]. Furthermore, it is well-established that FR dysfunction in humans, either through genetic mutation or autoantibodies blocking its function, causes birth defects and neurodevelopmental disorders in childhood [40–45], thus demonstrating that the role of FR in development differs between species. It is likely that in more complex biological systems, such as mammalian nervous system, FR-mediated folate endocytosis is required to maintain developmental processes, whereas the ubiquitous expression of FOLT-1 [22] is sufficient to ensure developmental integrity in *C. elegans*. Nevertheless, the lack of phenotypes in adult *folr-1* mutants highlights FR as an attractive and safe target for interventions that inhibit its function.

Although the role of FR in human and *C. elegans* development differs, its restricted expression is a common feature in both species. Interestingly, it has been shown that histone variant H2A.Z.1 and histone chaperone ASF1a promote the expression of FRα in cultured mouse neural precursor cells [46]. However, to our knowledge, it is not known which factors inhibit FR expression in a multicellular organism. Our data highlight the role of LIN-53/RbAp48 in restricting FORL-1 expression, thus raising the question of whether this histone-binding protein regulates FR expression also in humans. Tightly controlled expression of FR in both human and *C. elegans* implies that this receptor is detrimental for organismal health when expressed at high level. Indeed, we show that *folr-1* OE shortens lifespan in a microbiota- and microbial metabolism-dependent manner (Fig 6A, Fig 6D and Fig 6E).

Currently we do not know the mechanism of how elevated FOLR-1 level interacts with diet/microbiome to regulate lifespan. As mentioned earlier, FOLR-1 mediates the biological effects of bacterial folates [24], raising possibility that *folr-1* OE increases the uptake of these folates from the co-cultured bacteria, which then at high doses have adverse effect on health. Since N2 animals have shorter lifespan on OP50 compared to HT115 (Fig 6A) [5, 38], it is possible that HT115 bacteria produces less toxic bacterial folates. On the other hand, since *folr-1* OE shortens lifespan only on HT115 (Fig 6A), elevated FOLR-1 level could sensitize animals to HT115-produced bacterial folates, thus leading to shortened lifespan. Interestingly, metformin, a widely used drug to treat type 2 diabetes, promotes longevity on OP50 by disrupting bacterial folate and methionine cycles, whereas it does not extend lifespan on HT115 [5], which supports hypothesis that OP50 produces more toxic bacterial folates. Furthermore, we found that SMX-mediated inhibition of bacterial folate metabolism does not extend the lifespan of *folr-1* OE animals as much as with N2 (Fig 6D and Fig 6E), suggesting that overexpressed FOLR-1 can uptake bacterial folates from the pool that is generated even in the presence of SMX, which then suppresses SMX-induced longevity.

Regarding humans, the expression of FRs, and especially FRα, has been shown to be strongly increased in certain cancers [14, 15, 17–21]. Moreover, whereas RFC is considered as putative tumor suppressor, FRα is considered as putative oncogenic factor in human malignancies [47]. Notably, aberrant gut microbiota contributes to the onset and progression of several maladies, including cancer [48–50]. Although we cannot draw direct conclusions from *C. elegans* experiments to human cancer patients, our data raise the possibility that an interaction between an aberrant gut microbiota and increased FR expression may contribute to the systemic deterioration of health in cancer patients. Finally, although we are left with many open questions, this work highlights the importance of restricted FOLR-1 expression for both fitness and lifespan. Therefore, our data should encourage further studies on the role of this receptor in the organismal physiology, especially under conditions where it is overexpressed.

## Materials and Methods

### *C. elegans* strains and maintenance

For all experiments *C. elegans* were maintained on NGM plates (peptone, P4963, Merck; agar, A4550, Merck; NaCl, 746398, Merck) seeded with *E. coli* OP50 or HT115 bacteria carrying empty vector (EV, control vector for RNAi). The N2 (Bristol) strain was used as the wild-type. N2, GMC101 and CL2122 strains were obtained from the Caenorhabditis Genetics Center (CGC). *folr-1(syb4116)* X (PHX4116), *folr-1(syb5135)* X (PHX5135), *folr-1(syb4185)*[*folr-1::mNeonGreen*] X (PHX4185), *rps-6(syb7330[rps-6::wrmScarlet])* I (PHX7330) were created by using CRISPR/Cas9-mediated genome editing (SunyBiotech). All described crosses between genotypes were created within this study. *folr-1* OE strains (PHX4824 and PHX4825, *Is[Pfolr-1::folr-1::unc-54 3’UTR, Pmyo-2::gfp]*) were created by using microinjection (SunyBiotech). Transgenes were integrated into the genome by gamma irradiation (SunyBiotech). PHX4116 and PHX5135 strains were outcrossed two times with N2. PHX4824 and PHX4825 were outcrossed six times with N2. Related sequences for strains created within this study can be found from Supporting Information.

### RNA interference (RNAi)

*folt-1* RNAi was created by cloning part of *folt-1* ORF from *C. elegans* cDNA by using 5’-ataaccggtCCGTAAAGGAGTTTCGACCA-3’ and 5’-ataggtaccAAAATTGAAGCGACCAGTGC-3’ primers and ligating it into the L4440 vector. *lin-53* and *folr-1* RNAi clones were taken from Ahringer RNAi library, respectively. RNAi was performed by using the feeding protocol described earlier [51]. In *folr-1* RNAi experiments, animals were grown on RNAi for one generation (P0) before the experiment (qRT-PCR and lifespan analyses were performed with F1 generation).

### Imaging

For confocal imaging, *folr-1(syb4185)*[folr-1::mNeonGreen] X (PHX4185) animals were maintained at 20 °C and imaged at L3 larvae, L4 larvae and day 1 of adulthood. Animals were mounted on 2% agarose pads and immobilized by using 30 μM levamisole hydrochloride diluted in M9 solution. Leica SP8 upright confocal microscope was used for imaging. Z-stack images were acquired at 1.04 um slice intervals with HC PL APO 20x/0.75 IMM CORR CS2 objective. Z-stack images were converted to maximum projection-format by using Leica Application Suite X (LAS X) software. For *lin-53* RNAi experiment, *folr-1(syb4185)*[folr-1::mNeonGreen] X (PHX4185) animals were grown on RNAi from hatching at 20 °C. L4 larvae animals were mounted on 2% agarose pads and immobilized by using 30 μM levamisole hydrochloride diluted in M9 solution. Imaging was done with Olympus BX63 microscope by using 10x objective. Similarly, *rps-6(syb7330[rps-6::wrmScarlet])* I (PHX7330) animals were mounted on 2% agarose pads as L4 larvae and immobilized by using 30 μM levamisole hydrochloride diluted in M9 solution. Imaging was done with Olympus BX63 microscope by using 10x objective.

### Lifespan analysis

Except for one experiment done at 25 degrees Celsius (°C), all *C. elegans* lifespan experiments were done at 20 °C. Lifespan experiments were initiated by letting gravid hermaphrodites (P0 generation) to lay eggs on NGM agar plates, and F1 generation was scored for lifespan. Alternatively, animals were bleached and let to hatch overnight in M9 before plating L1 larvae to experimental plates. These two alternative ways to initiate lifespan did not affect the conclusions made from the experiments. For folate supplementation experiments, folic acid (FA, Merck, #F8758) and 5-methyltetrahydrofolate (5-MTHF, Merck, #M0132) were added to NMG media at indicated concentrations. Sulfamethoxazole (SMX, Merck, #S7507) was added to NMG media at final concentration of 128 μg/ml [6, 7]. At L4 larval stage animals were transferred to plates containing 10 µM of 5-Fluorouracil (Merck, #F6627) to prevent progeny production. Animals that had exploded vulva or that crawled off the plate were censored. Animals were counted as dead if there was no movement after poking with a platinum wire. Lifespans were checked every 1-3 days. Mean lifespan ± standard error (SE) is reported in S1 Table.

### Activity measurement

Animals were synchronized by bleaching and plated as L1 larvae on NGM agar plates, which were put to 20 °C. At L4 larval stage animals were transferred to plates containing 10 µM of 5-Fluorouracil (Merck, #F6627) to prevent progeny production. The activity of N2, *folr-1* mutants and *folr-1* OE strains was measured on day 4 of adulthood. GMC101 and CL2122 (and crosses of these strains with *folr-1(sy4116)* mutant) were transferred to 25 °C at day 1 of adulthood, and the activity was measured at day 2 of adulthood. For activity measurement, 10 animals were put to single well of 96-well plate containing 100 μl of M9 solution. 10-12 wells were used per experiment for each strain. Activity was measured for two hours with wMicroTracker (InVivo Biosystems).

### Progeny count

Single L4 stage N2 and *folr-1(syb4116)* mutants were placed on 10 small agar plates. Animals were transferred to new plates daily during the reproduction period, and viable offspring were counted.

### RNA-seq

N2 and *folr-1(syb4116)* animals were synchronized by bleaching and plated as L1 larvae on NGM agar plates. Animals were collected at L4 larval stage (three biological replicates for both strains) and frozen in liquid nitrogen. Total RNA was extracted with TRIzol Reagent (ThermoFisher Scientific, #15596018). Samples were sent to Novogene for library construction, quality control and sequencing. In short, mRNA was purified from total RNA using poly-T oligo-attached magnetic beads. After fragmentation, the first strand cDNA was synthesized using random hexamer primers, followed by the second strand cDNA synthesis using either dUTP for directional library or dTTP for non-directional library. The library preparations were sequenced on an Illumina platform and paired-end reads were generated. To obtain clean reads, raw data (raw reads) of FASTQ format were firstly processed through fastp. Paired-end clean reads were mapped to the reference genome using HISAT2 software. FeatureCounts was used to count the read numbers mapped of each gene. Consequently, RPKM of each gene was calculated based on the length of the gene and reads count mapped to this gene. Prior to differential gene expression analysis, for each sequenced library, the read counts were adjusted by Trimmed Mean of M-values (TMM) through one scaling normalized factor. Differential expression analysis between two conditions (three biological replicates per condition) was performed using DESeq2 R package [52]. The resulting P values were adjusted using the Benjamini and Hochberg’s approach for controlling the False Discovery Rate (FDR). Genes with an adjusted P value < 0.05 found by DESeq2 were assigned as differentially expressed. Differential expression analysis of two conditions was performed using the edgeR R package [53]. The P values were adjusted using the Benjamini and Hochberg methods. Corrected pvalue of 0.005 and |log2^(Fold Change)^| of 1 were set as the threshold for significantly differential expression. KEGG pathway enrichment analysis was done with clusterProfiler R package. The RNA-seq data are available in the Gene Expression Omnibus (GEO) database repository (GSE227272).

### Quantitative RT-PCR (qRT-PCR)

Animals were synchronized by bleaching and plated as L1 larvae on NGM agar plates, which were put to 20 °C. Animals were collected at L4 stage and frozen in liquid nitrogen. TRIzol Reagent (ThermoFisher Scientific, #15596018) was used to extract RNA. cDNA synthesis was done with QuantiTect Reverse Transcription Kit (Qiagen, #205313) and qRT-PCR reactions were run with HOT FIREPol SolisGreen qPCR Mix-reagent (Solis BioDyne, #08-46-00001) using CFX384 machine (Bio-Rad). qRT-PCR data was normalized to the expression of *cdc-42* and *pmp-3*. qRT-PCR oligos used in this study are provided in S2 Table. qRT-PCR experiments were performed with three biological replicates. Analysis of ribosomal subunit expression was performed once with each *folr-1* mutant, and the analysis of *folr-1* RNAi efficiency in N2 and *folr-1* OE strains was performed twice with similar results. Statistical significances were analyzed by using Student’s t-test and one-way ANOVA.

### Western blot

Animals were synchronized by bleaching and plated as L1 larvae on NGM agar plates, which were placed in 20 °C. At L4 larval stage, animals were transferred to plates containing 10 µM of 5-Fluorouracil (Merck, #F6627) to prevent progeny production. N2, and *folr-1(syb4116)* mutants were collected on day 2 of adulthood and frozen in liquid nitrogen. GMC101 and CL2122 (and crosses of these strains with *folr-1(sy4116)* mutant) were transferred to 25 °C at day 1 of adulthood, collected at day 2 of adulthood, and frozen in liquid nitrogen. Animals were lysed in protease inhibitor cocktail (ThermoFisher Scientific, #78430)-supplemented urea solution (Merck, #51457) by grinding with plastic pestle in 1.5 ml Eppendorf tubes. Lysates were resolved on 4–15% precast polyacrylamide gels (Bio-Rad, #4561083). Immun-Blot PVDF Membrane (Bio-Rad, #1620177) was used for blotting. Purified anti-β-Amyloid, 1-16 Antibody (6E10) (used with 1:1000 dilution) was purchased from BioLegend. Anti-S6 Ribosomal Protein Antibody (used with 1:1000 dilution) was purchased from Merck (#ZRB1172). α-tubulin antibody (used with 1:5000 dilution) was purchased from Merck (#T5168). Clarity Western ECL Substrate (Bio-Rad, #1705061) and ChemiDoc MP-imager (Bio-Rad) were used for protein detection in Western blot.

### Statistical analysis

Statistical analyses for qRT-PCR data were carried out in GraphPad Prism and Excel, and the data represent the mean of three biological replicates ± standard deviation (SD). Statistical analyses for activity measurements and Western blot data were carried out in GraphPad Prism and Excel, respectively. Statistical details can be found in the figures and figure legends. Statistical analyses for lifespan experiments were carried out in R by using the Cox-proportional hazard regression. Statistical details for the lifespan data can be found in S1 Table.

## Acknowledgements

The authors thank Drs Susana Garcia and Carina Holmberg (University of Helsinki) for sharing reagents. Some strains were provided by the CGC, which is funded by NIH Office of Research Infrastructure Programs (P40 OD010440). This work was supported by Research Council of Finland and University of Helsinki.

## Supporting information captions

**S1 Fig. Confocal images of L3 larvae and day 1 adult N2 (wild-type) and FOLR-1::mNeonGreen-expressing animals.**

N2 is used as a control to recognize background signal. Arrows indicate FOLR-1::mNeonGreen localization.

**S2 Fig. Lifespan upon heat stress and 250 mM folic acid (FA) treatment.**

(A) Lifespan of N2 and *folr-1(syb4116)* mutants at 25 degrees Celsius (25 °C). (B) Lifespan of N2 animals on plates supplemented with 250 mM folic acid (FA). Lifespan statistics are reported in S1 Table.

**S3 Fig. The expression of ribosome subunits in *folr-1* mutants.**

(A) qRT-PCR of selected ribosome subunits in L4 stage *folr-1(syb4116)* and (B) *folr-1(syb5135)* mutants compared to N2. Bars represent mRNA levels relative to N2 with error bars indicating mean ± SD of three biological replicates, each with three technical replicates (*p < 0.05, **p < 0.01, unpaired Student’s t-test). (C) Representative images of L4 stage animals expressing RPS-6::wrmScarlet-fusion protein in wild-type- and *folr-1(syb5135)* background. (D) Representative Western blot of ribosomal RPS-6 in day 2 adult N2 and *folr-1(syb4116)* mutants. Blot shows three technical replicates from one biological replicate. (E) Quantification of RPS-6 Western blots of day 2 adult N2 and *folr-1(syb5135)* mutants. Bars represent the fold change of tubulin-normalized RPS-6 level relative to N2 with error bars indicating mean ± SD of three biological replicates (statistical significance calculated with unpaired Student’s t-test).

